# Design-of-Experiments In Vitro Transcription Yield Optimization of Self-Amplifying RNA

**DOI:** 10.1101/2021.01.08.425833

**Authors:** Karnyart Samnuan, Anna K. Blakney, Paul F. McKay, Robin J. Shattock

**Author notes:** Corresponding Author: Robin J. Shattock.

## Abstract

Self-amplifying RNA (saRNA) vaccines are able to induce a higher antigen-specific immune response with a more cost-effective and rapid production process compared to plasmid DNA vaccines. saRNAs are synthesized through in vitro transcription (IVT) however; this process has mainly been optimized for relatively short mRNAs. Here, we optimized the IVT process for long saRNAs, approximately 9.4 kb through a design of experiment (DoE) approach to produce a maximal RNA yield and validated the optimal IVT method on various sizes of RNA. We found that magnesium has the highest impact on RNA yield with acetate ions enabling a higher yield than chloride ions. In addition, the interaction between magnesium and nucleoside triphosphates (NTPs) is highly essential for IVT. Further addition of sodium acetate (NaOAc) during IVT provided no added benefit in RNA yield. Moreover, pyrophosphatase was not essential for productive IVT. The optimal IVT method can be used to synthesize different lengths of RNA. These findings emphasize the ability to synthesize high quality and quantity of saRNA through IVT and that the optimal amount of each component is essential for their interactions to produce a high RNA yield.

## Introduction

The use of RNA-based vaccines as a vaccine platform against infectious diseases have emerged over the past decade because of advances in RNA production and formulation. RNA vaccines can trigger stronger and more potent cellular and humoral immune response than plasmid DNA (pDNA) and avoid any potential risk of host cell genome integration^1^. Self-amplifying RNA (saRNA) vaccines in particular, are advantageous in vaccine research due to their ability to self-replicate, allowing exponential expression of antigen^2^. Compared to the conventional mRNA vaccination approach, saRNA can induce equivalent protection with lower doses^3^ and has also been shown to elicit longer antigen expression *in vivo*^4^. Furthermore, stronger cellular and humoral responses are induced when mice are immunized with saRNA compared to regular mRNA^5^. Over the past two decades, many *in vivo* studies have shown that saRNA is able to induce potent and robust protection against infectious diseases like HIV-1^6, 7^, influenza virus^3, 8, 9^, dengue virus^10, 11^, respiratory syncytial virus (RSV)^9, 12^ and Ebola virus^13, 14^.

saRNA, or an RNA replicon, is a single-stranded RNA derived from a positive strand virus genome, such as alphaviruses^15^. The non-structural proteins of the virus encode for a replicase, an enzyme complex that catalyzes the self-replication of RNA template, producing a high number of RNA copies^15^. The viral structural proteins, which are downstream of the non-structural proteins, are replaced with a gene(s) of interest (GOI), preventing the formation of an infectious viral particle. RNA replicons are produced by in vitro transcription (IVT) of linearized DNA templates using a T7 RNA polymerase that binds to the T7 promoter of the template to read off the sequence and synthesize RNA molecules^16^. A typical IVT reaction for the synthesis of RNA includes: i) a linearized pDNA with a T7 promoter; ii) nucleoside triphosphates (NTPs) for the four bases; iii) a ribonuclease inhibitor for inactivating RNase; iv) a pyrophosphatase for degrading accumulated pyrophosphate; v) magnesium anion which is a cofactor for the T7 polymerase; vi) a pH buffer that contains optimal concentrations of an antioxidant and a polyamine^17^.

Contemporary IVT protocols have been primarily developed and optimized for small RNAs (<100 nt)^18, 19^, but not for saRNAs. Because saRNAs are much longer and have a high degree of secondary structure, the optimal conditions for IVT may be different than for mRNA. Therefore, we aim to use a design of experiment (DoE) approach to optimize the production of long RNA replicons (approx. 9.4 kb) through IVT. DoE is a method used to determine the relationship between each factor or variable that are known to influence a particular process or an output of a process^20^. By using this approach, we sought to establish a mathematical relationship between each variable as well as determine the most influential factor in order to maximize the output.

Here, we used a variety of designs such as full factorial designs and definitive screening designs to understand the relationship between each component that are thought to be necessary in synthesizing RNA replicons through IVT as shown in Figure 1. Firstly, a full two-level factorial DoE was conducted to determine which components have the highest significance in IVT as well as their secondary interactions. A DoE with a low, middle and high point for all of the components would not be feasible as that would result in >8000 (20^3^ reactions). Hence, we used a factorial design to screen the initial components. A full three-level factorial design was then utilized to explore the relationship between sodium (Na^+^), magnesium (Mg^+2^) and NTPs as well as to determine the difference between acetate ions and chloride ions in order to yield high RNA concentrations. Next, we titrated the concentration of magnesium acetate (MgOAc_2_) to generate a definitive optimal ratio between NTPs and Mg^+2^. We also titrated the T7 RNA polymerase concentration and subsequently increased the sodium acetate (NaOAc) concentration at different timepoints during IVT to see whether this can increase RNA yield. Once the initial components were screened, we used a definitive screening design to optimize and analyze curvature of the design space. Here, a definitive screening design was then performed to understand the significance of incorporating pyrophosphatase, spermidine, Dimethyl sulfoxide (DMSO), betaine and a surfactant of either Triton X-100 or Tween 20. Lastly, we made different sizes of RNA using the optimal IVT method and transfected these RNAs *in vitro* before performing flow cytometry to confirm protein expression.

**Figure 1.**
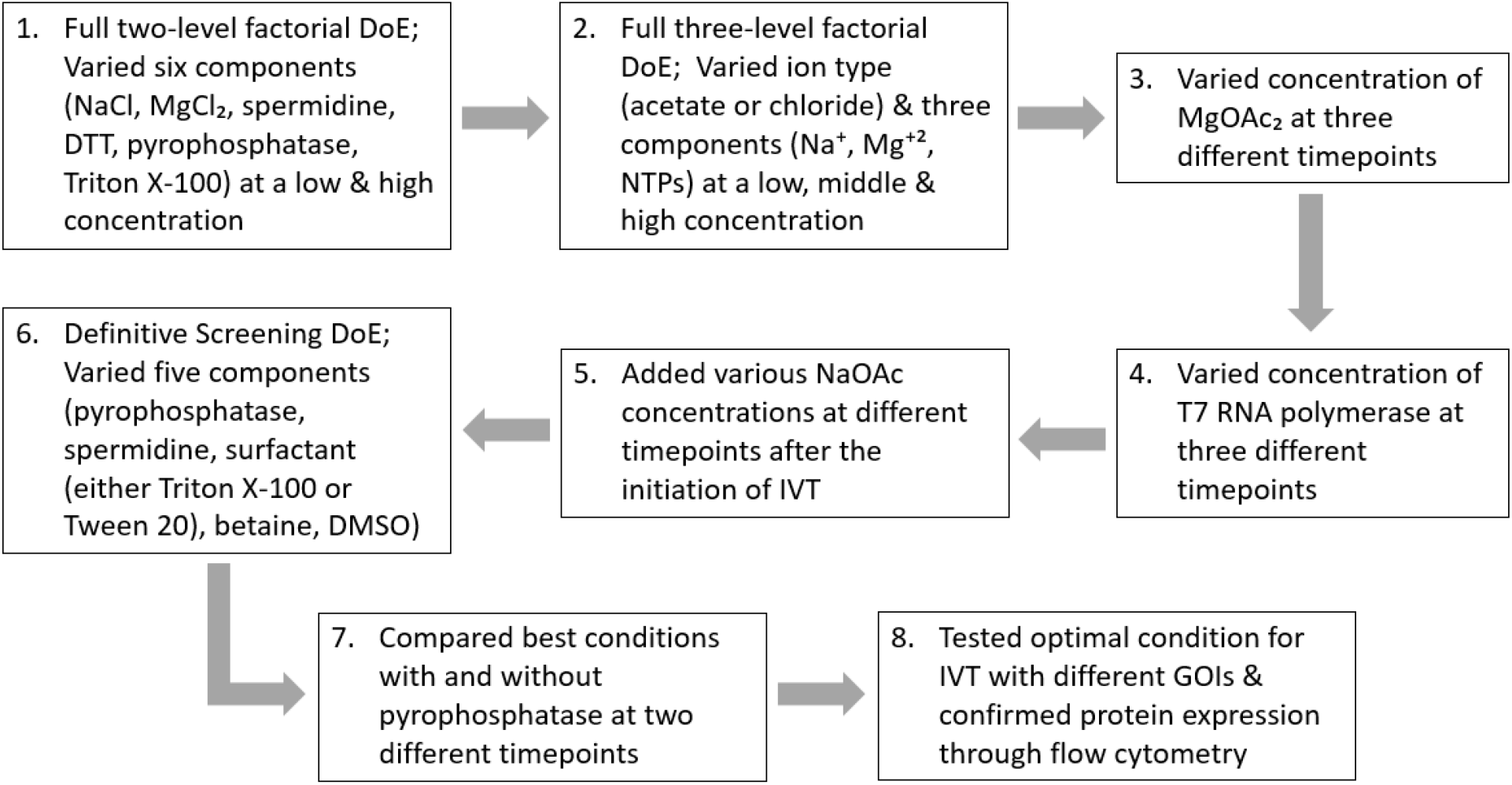
A visual flow diagram of all the DoEs performed for this study in order to achieve our optimal condition for IVT.

## Materials and Methods

### Plasmid DNA Synthesis and Purification

The plasmid DNA (pDNA) construct used for the synthesis of RNA replicons encodes for the non-structural proteins of Venezuelan Equine Encephalitis Virus (VEEV), wherein firefly luciferase (fLuc) gene (GenBank: AB762768.1) (GeneArt, Germany) was cloned into the plasmid right after the sub-genomic promoter using the restriction sites NdeI and MluI-HF. The pDNA was transformed into Escherichia coli and grown in 100 mL cultures in lysogeny broth (LB) media with 100 μg/mL carbenicillin (Sigma-Aldrich, U.S). Isolation and purification of the pDNA was done using a Plasmid Plus maxi kit (QIAGEN, UK) and a NanoDrop One Microvolume UV-Vis Spectrophotometer (ThermoFisher, UK) was used to measure the concentration and purity of the pDNA. The pDNA sequence was confirmed with Sanger sequencing (GATC Biotech, Germany). Prior to RNA IVT, pDNA was linearized using MluI for 3h at 37 °C, according to the manufacturer’s instructions.

### RNA Synthesis and Quantification

Each of the components listed in Table 1 were used for certain DoEs. 1 μg of linearized DNA template was used and kept constant across all IVT reactions with a final volume of 100 μL per reaction which were incubated at 37 °C for either 2, 4 or 6 h. RNA yield was measured right after IVT using the Qubit RNA Broad Range Assay kit with the Qubit Fluorometer (Thermo Fisher, UK) according to the manufacturer’s protocol. Each variable of each DoE was performed in triplicate.

**Table 1.** The list of components used for the different DoEs.

| **Components used for IVT:** | **Original Stock Concentration**<br>**for Each Component:** | **Manufacturer:** |
| --- | --- | --- |
| **1 µg DNA template** | - | - |
| **ATP, CTP, UTP and GTP** | 100 mM | Thermo Fisher, UK |
| **Tris Hydrochloride (Tris HCl) pH= 8** | 1000 mM | Thermo Fisher, UK |
| **HEPES pH= 7.0-7.6** | 1000 mM | Sigma-Aldrich, US |
| **RNase Inhibitor** | 40 U/ µL | Thermo Fisher, UK |
| **T7 RNA polymerase** | 50 U/ μL | Thermo Fisher, UK |
| **Sodium Chloride (NaCl)** | 5000 mM | Thermo Fisher, UK |
| **Magnesium Chloride (MgCl<sub>2</sub>)** | 1000 mM | Thermo Fisher, UK |
| **Spermidine** | 100 mM | Sigma-Aldrich, US |
| **Dithiothreitol (DTT)** | 1000 mM | AppliChem, Germany |
| **Pyrophosphatase** | 0.1 U/ μL | Thermo Fisher, UK |
| **Triton X-100** | 100 X | Sigma-Aldrich, US |
| **Sodium Acetate (NaOAc)** | 3000 mM | Sigma-Aldrich, US |
| **Magnesium Acetate (MgOAc<sub>2</sub>)** | 1000 mM | Sigma-Aldrich, US |
| **Tween 20** | 10 % | Thermo Fisher, UK |
| **Betaine** | 5 M | Sigma-Aldrich, US |
| **Dimethyl sulfoxide (DMSO)** | 100 % | Sigma-Aldrich, US |

### Design of Experiment and Statistical Analysis

JMP software, version 14.1 was used to carry out all the DoE analysis using mainly full factorial designs and one definitive screening design with RNA yield as the response for each experiment. A fit model of standard least squares for effect screening was used to analyze the data of each DoE, where factors that were found to be non-significant were removed in order to improve the models and refine the optimal factors for IVT^21^. Graphs were prepared using GraphPad Prism, version 8.2.

### RNA Purification and RNA Gel

After IVT, RNA was purified using MEGAclear™ Transcription Clean-up Kit (Thermo Fisher, UK) according to the manufacturer’s protocol. In order to access the quality of the RNA, purified RNAs and the RNA Millennium Marker Ladder (Thermo Fisher, UK) were mixed with 2x RNA loading dye (Thermo Fisher, UK) and incubated at 50 °C for 30 min to denature the RNA. A 1.2 % agarose gel with 1x NorthernMax Running Buffer (Thermo Fisher, UK) was prepared. After incubation, the denatured ladder and samples were added to the gel and the gel was ran at 80 V for 45 min. The gel was then imaged on a GelDoc-It2 (UVP, UK).

### Cells and *In Vitro* Transfections

HEK293T.17 cells (ATCC, USA) were cultured in complete Dulbecco’s Modified Eagle’s Medium (DMEM) (Gibco, Thermo Fisher, UK) containing 10 % fetal bovine serum (FBS), 1 % L-glutamine and 1 % penicillin-streptomycin (Thermo Fisher, UK). Cells were plated in a 6-well plate at a density of 1.08 × 10^6^ cells per well 48 h prior to transfection. Transfection of saRNAs encoding different GOIs and mRNA fLuc was performed using Lipofectamine MessengerMAX (Thermo Fisher, UK) according to the manufacturer’s instructions.

### Flow Cytometry

Transfected cells were harvested and resuspended in 1mL of FACS buffer (PBS + 2.5 % FBS) at a concentration of 1 x 10^7^ cells /mL. 100 μL of the resuspended cells was added to a FACS tube and stained with 50 μL of Live/Dead Fixable Aqua Dead Cell Stain (Thermo Fisher, UK) at a 1:400 dilution on ice for 20 min. Cells were then washed with 2.5 mL of FACS buffer and centrifuged at 1750 rpm for 7 min. The cells that were transfected with a fLuc RNAs were permeabilized with Fixation/ Permeabilization solution kit (BD Biosciences, UK) for 20 min before washing them with 2.5 mL of FACS buffer and centrifuging at 1750 rpm for 7 min. Cells transfected with the fLuc RNAs were stained with 5 μL of anti-Luciferase antibody (C-12) PE: sc-74548 PE (Santa Cruz Biotechnology, US) while the MDR1 replicons were stained with 5 μL of PE anti-human CD243 (ABCB1) antibody clone 4E3.16 (Biolegend, US). After staining for 30 min on ice, cells were washed with 2.5 mL of FACS buffer, centrifuged at 1750 rpm for 7 min and resuspended with 250 μL of PBS. Cells were fixed with 250 μL of 3 % paraformaldehyde for a final concentration of 1.5 %. Samples were analyzed on a LSRForterssa (BD Biosciences, UK) with FACSDiva software (BD Biosciences, UK). Data were analyzed using FlowJo Version 10 (FlowJo LLC, Ashland, OR, USA).

## Results

### Magnesium is the most significant component for influencing IVT yield of saRNA

A full factorial design of experiment was designed with six factors, including NaCl, MgCl_2_, spermidine, Dithiothreitol (DTT), pyrophosphatase and Triton X-100, which were varied at a low and high concentration (Table 2). The RNA yield quantified were entered to the JMP software and a fit model of standard least squares was generated.

**Table 2.** The components used for a full two-level factorial DoE.

| **Fixed components** | **Concentration of fixed components** | **Components that were varied** | **Concentrations of components varied** |
| --- | --- | --- | --- |
| Linearized DNA Template | 1 µg | Sodium Chloride (NaCl) | 2.5 mM |
|  |  |  | 25 mM |
| ATP, CTP, UTP and GTP | 4 mM each | Magnesium Chloride (MgCl<sub>2</sub>) | 2.4 mM |
|  |  |  | 24 mM |
| Tris Hydrochloride (Tris HCl) | 40 mM | Spermidine | 0.2 mM |
|  |  |  | 2 mM |
| RNase Inhibitor | 4 U | Dithiothreitol (DTT) | 4 mM |
|  |  |  | 40 mM |
| T7 RNA Polymerase | 100 U | Pyrophosphatase | 0.01 U |
|  |  |  | 0.1 U |
|  |  | Triton X-100 | 0 % |
|  |  |  | 0.01 % |

MgCl_2_ had the highest significance in IVT with a log worth of 73.862. Spermidine was the second most significant component for IVT with a log worth of 4.070 and its interactions with NaCl and MgCl_2_ were also statistically significant for enhancing high RNA yield. However, DTT, pyrophosphatase and Triton X-100 all showed negligible impact in IVT (p= 0.20622, 0.56417, and 0.66715, respectively), (Table 3 and Figure 2).

**Table 3.** The significance of each component varied and their interactions. A log worth > 2 or a p-value < 0.01 is considered significant.

| Source | Log Worth | P-value |
| --- | --- | --- |
| $\text{MgCl}_2$ (2.4, 24) | 73.862 | 0.00000 |
| $\text{NaCl}^*\text{Spermidine}$ | 8.144 | 0.00000 |
| Spermidine (0.2, 2) | 4.070 | 0.00009 |
| $\text{MgCl}_2^*\text{Spermidine}$ | 4.070 | 0.00009 |
| $\text{NaCl}^*\text{DTT}$ | 2.536 | 0.00291 |
| $\text{NaCl}$ (2.5, 25) | 1.857 | 0.01390 |
| $\text{NaCl}^* \text{MgCl}_2$ | 1.857 | 0.01390 |
| $\text{NaCl}^*\text{Triton X-100}$ | 1.631 | 0.02340 |
| $\text{DTT}^*\text{Triton X-100}$ | 1.391 | 0.04060 |
| Spermidine*DTT | 1.233 | 0.05848 |
| DTT (4, 40) | 0.686 | 0.20622 |
| Pyrophosphatase (0.01, 0.1) | 0.249 | 0.56417 |
| Triton X-100 (0, 0.01) | 0.176 | 0.66715 |

**Figure 2.**
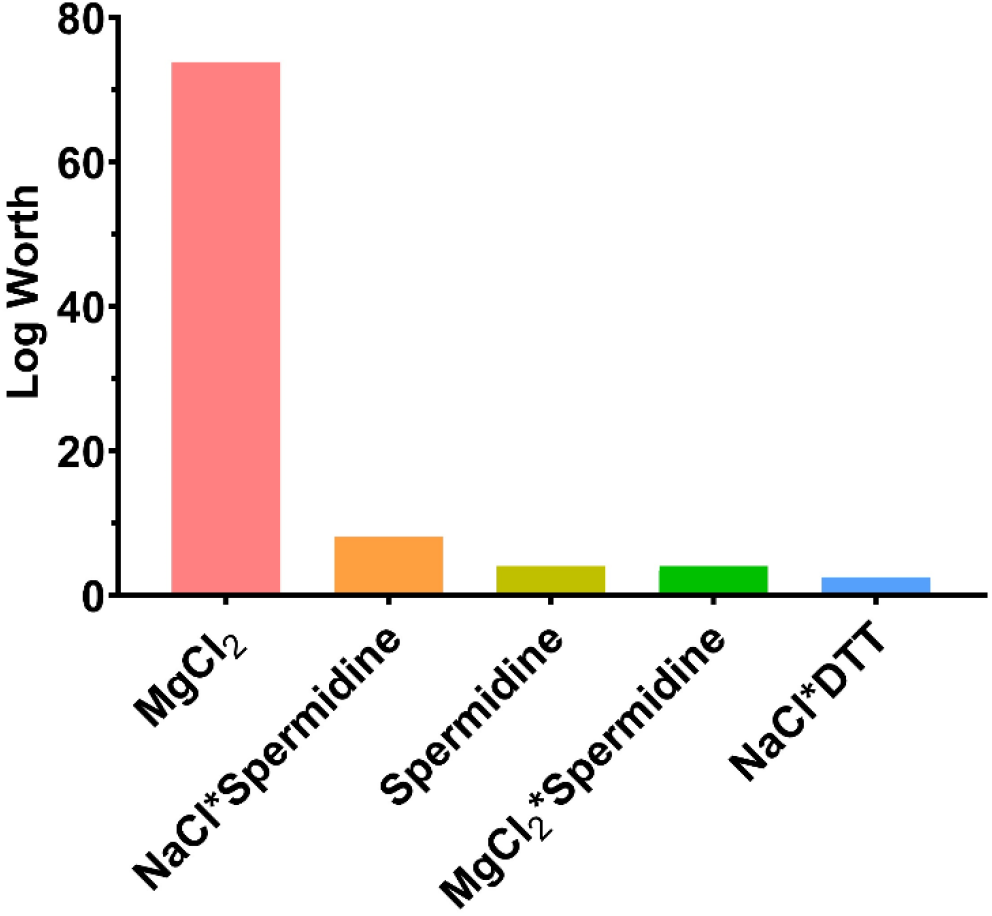
The most significant components varied based on a full two-level factorial DoE analysis. A log worth > 2 is considered significant. Each variable of the DoE was done in triplicates.

Furthermore, MgCl_2_ also had the highest coefficient estimates compared to other components and its secondary interactions (Figure 3).

**Figure 3.**
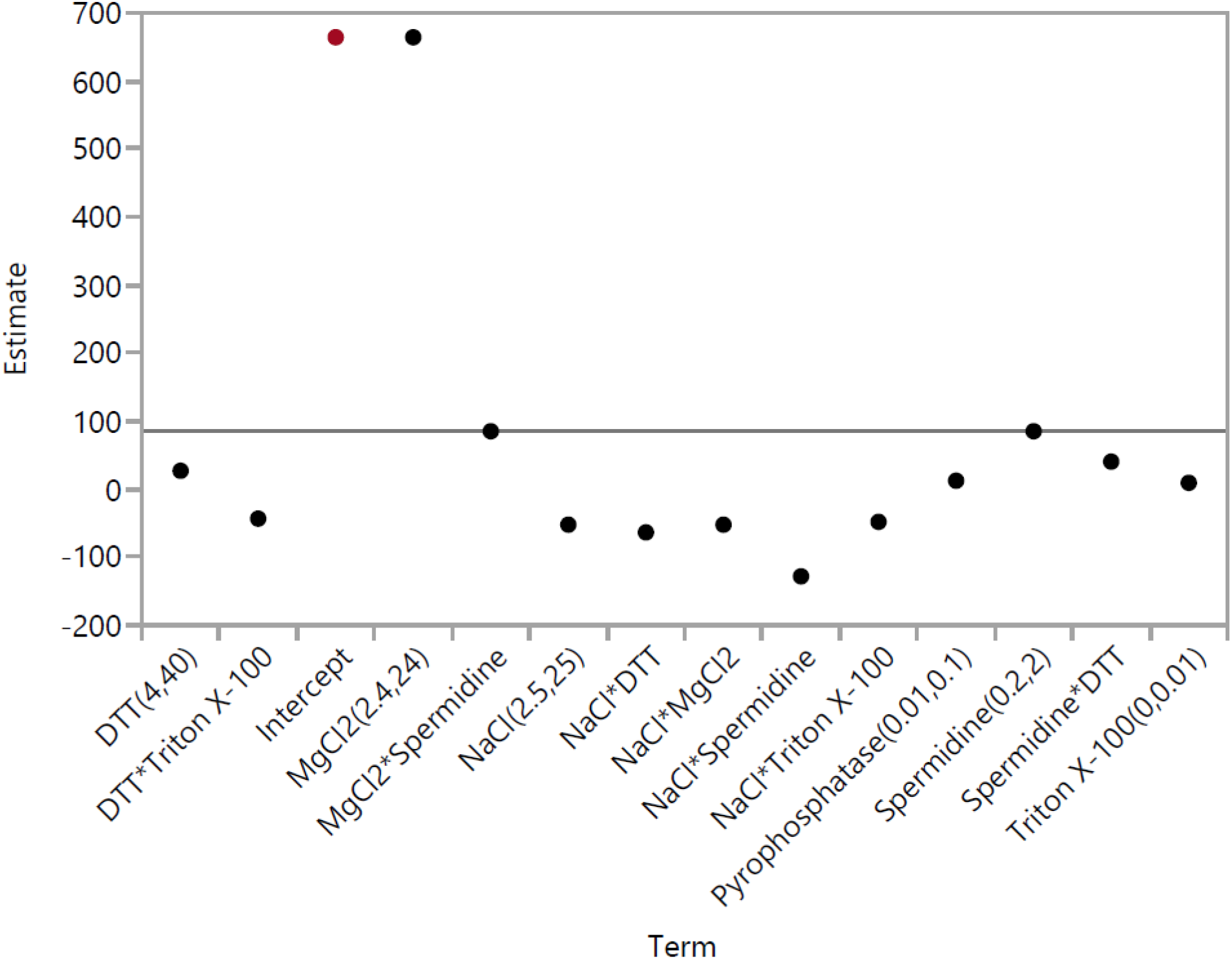
The coefficient estimates of each component and their secondary interactions based on a full two-level factorial DoE analysis. Each component’s coefficient estimate corresponds to the change in the mean response for each level and the average response across all levels.

The model equation showing the mathematical relationship between each variable in order to give a specific RNA yield is:

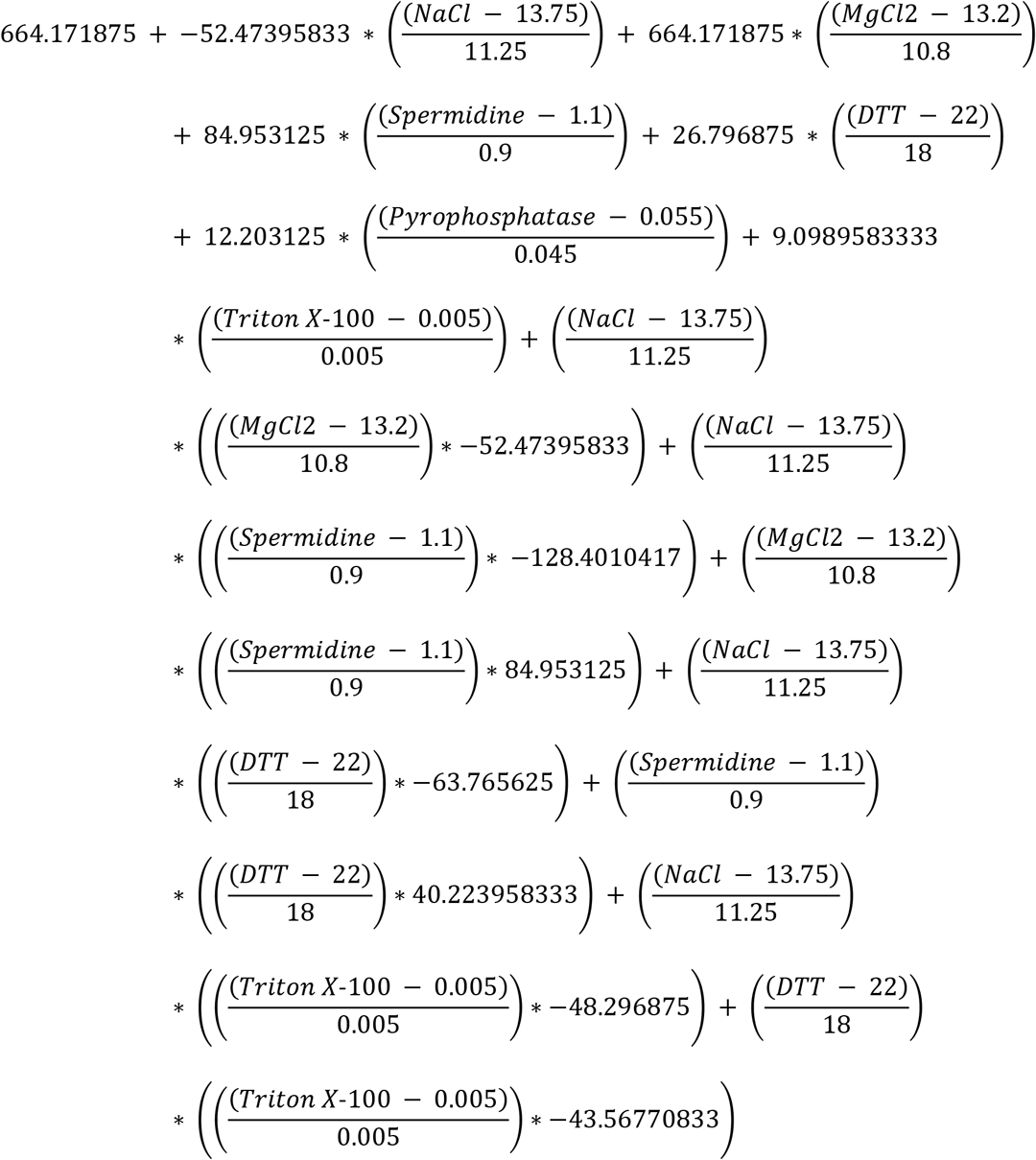

### Acetate ions are more effective than chloride ions in enhancing RNA yield

We then sought to examine the interactions between the concentrations of magnesium (Mg^+2^), sodium (Na^+^) and NTPs as well as the significance of varying the ion type between acetate and chloride. Hence, a full three-level factorial design of experiment was performed with three factors (Mg^+2^, Na^+^, NTPs) varied at three different concentrations (Table 4). Because DTT showed no significance in the previous full two-level factorial DoE but had an estimate of more than zero, we decided to use 40 mM DTT as the fixed component for this DoE. Furthermore, the previous DoE showed that spermidine mainly had significant secondary interactions with NaCl and MgCl_2_ but because the main focus of this DoE is to better understand the main interactions between Na^+^, Mg^+2^ and NTPs, we decided to use 0.2 mM spermidine as the fixed component here.

**Table 4.**
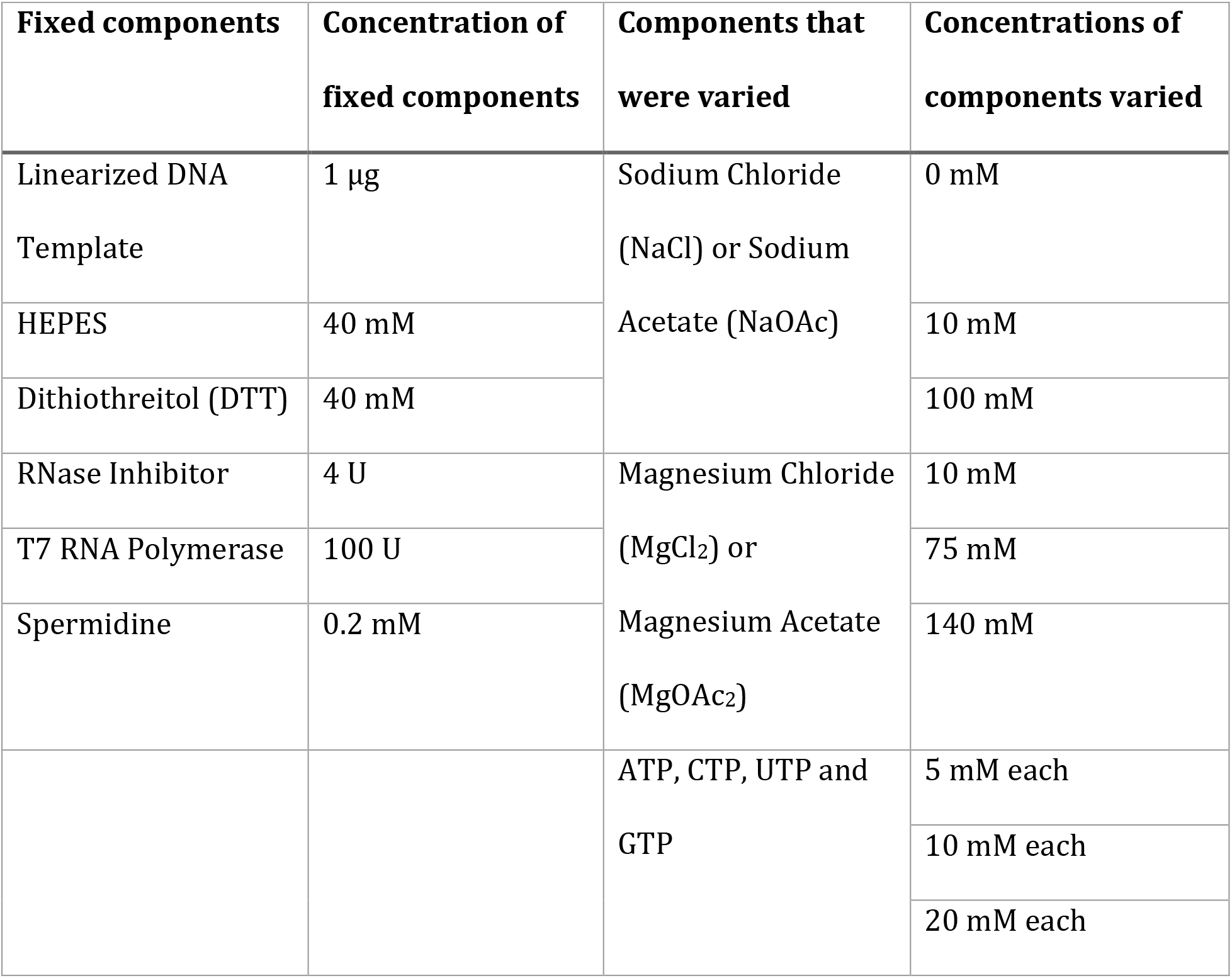
The components used for a full three-level factorial DoE.

| Fixed components | Concentration of fixed components | Components that were varied | Concentrations of components varied |
| --- | --- | --- | --- |
| Linearized DNA Template | 1 $\mu\text{g}$ | Sodium Chloride (NaCl) or Sodium Acetate (NaOAc) | 0 mM |
| HEPES | 40 mM |  | 10 mM |
| Dithiothreitol (DTT) | 40 mM |  | 100 mM |
| RNase Inhibitor | 4 U | Magnesium Chloride ($\text{MgCl}_2$) or Magnesium Acetate ($\text{MgOAc}_2$) | 10 mM |
| T7 RNA Polymerase | 100 U |  | 75 mM |
| Spermidine | 0.2 mM |  | 140 mM |
|  |  | ATP, CTP, UTP and GTP | 5 mM each |
|  |  |  | 10 mM each |
|  |  |  | 20 mM each |

We found that the ion type plays a significant role in IVT (Table 5 and Figure 4A) with acetate ions being more effective than chloride ions (p= 0.00906) (Figure 4B). The coefficient estimates graph also showed that acetate ions have the highest estimate (Figure 5). Further examination of interaction between Mg^+2^ and NTPs indicated that the highest RNA yield was achieved with a Mg^+2^ concentration of 75 mM and NTPs concentration of 10 mM each (Figure 4C). Further increasing the concentration of each NTP had a deleterious effect on IVT, causing RNA yield to plateau. These data suggest the optimal balance of Mg^+2^ and NTPs needed for effective IVT and maximal RNA quantity requires a molecular ratio of 1: 1.875 between total NTPs and Mg^+2^.

**Table 5.** The significance of each component varied. A log worth > 2 or a p-value < 0.01 is considered significant.

| Source | Log Worth | P-value |
| --- | --- | --- |
| Ion Type | 2.043 | 0.00906 |
| NTPs | 1.898 | 0.01265 |
| $Mg^{+2}$ | 1.207 | 0.06212 |
| $Na^{+}$ | 0.448 | 0.35657 |

**Figure 4.**
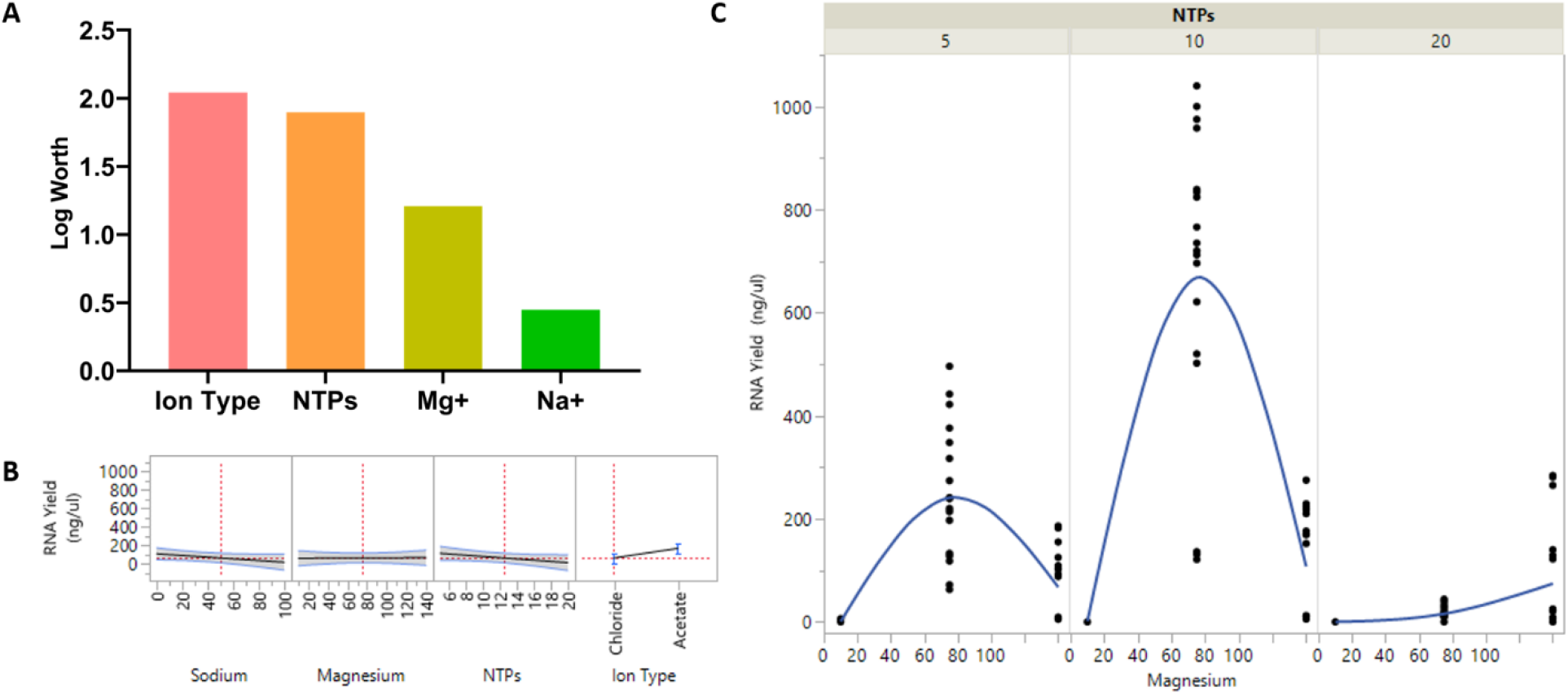
A full three-level factorial DoE analysis of saRNA synthesis through IVT. (A) The significance of each component varied represented in a graph; (B) The effects of each component varied on the RNA yield is plotted next to each other to understand the relationship between each component in order to produce high RNA yield; (C) The relationship between NTPs and Mg^+2^ at different concentrations are shown in relation to the RNA yield. Each variable of the DoE was done in triplicates. A log worth > 2 is considered significant.

**Figure 5.**
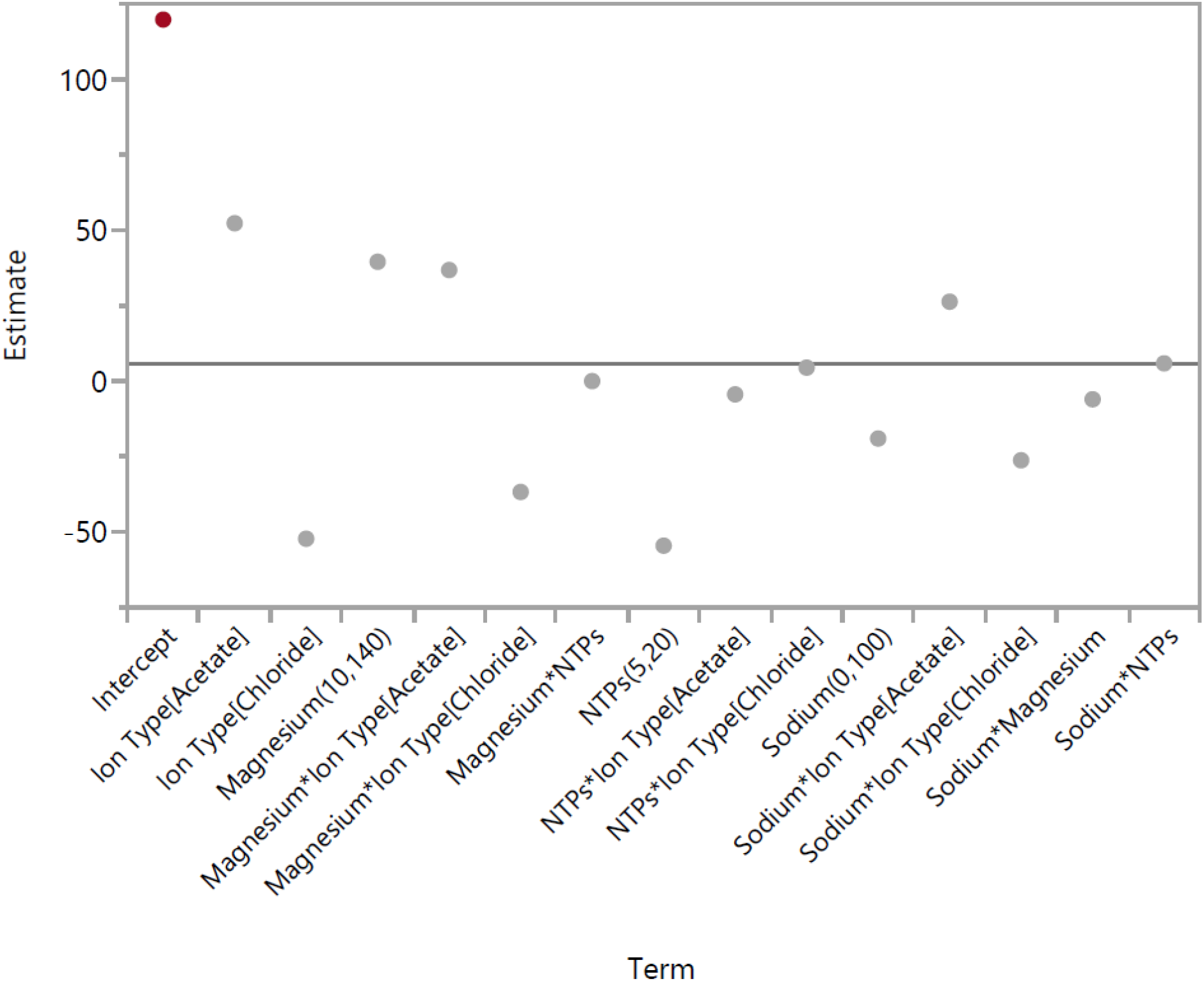
The coefficient estimates of each component and their secondary interactions based on a full three-level factorial DoE analysis. Each component’s coefficient estimate corresponds to the change in the mean response for each level and the average response across all levels.

### 75mM MgOAc_2_ is the optimal concentration for all incubation periods

Based on the observation that Mg^+2^ is the most significant component for IVT, and that acetate ions result in higher yield than chloride ions, we wanted to examine the optimal concentration of MgOAc_2_ and incubation time in order to produce maximal RNA yield.

The concentration of NTPs were kept at 10 mM each while varying the concentrations of MgOAc_2_ ranging from 25 - 125 mM. RNA yields were measured after 2, 4 and 6 h of incubation. As shown in Figure 6, the general trend shown for all timepoints is that the RNA yield reaches its highest at 75mM MgOAc_2_ and then it slowly decreases with higher concentrations of MgOAc_2_. In addition, the longer the incubation period, the higher the RNA yield.

**Figure 6.**
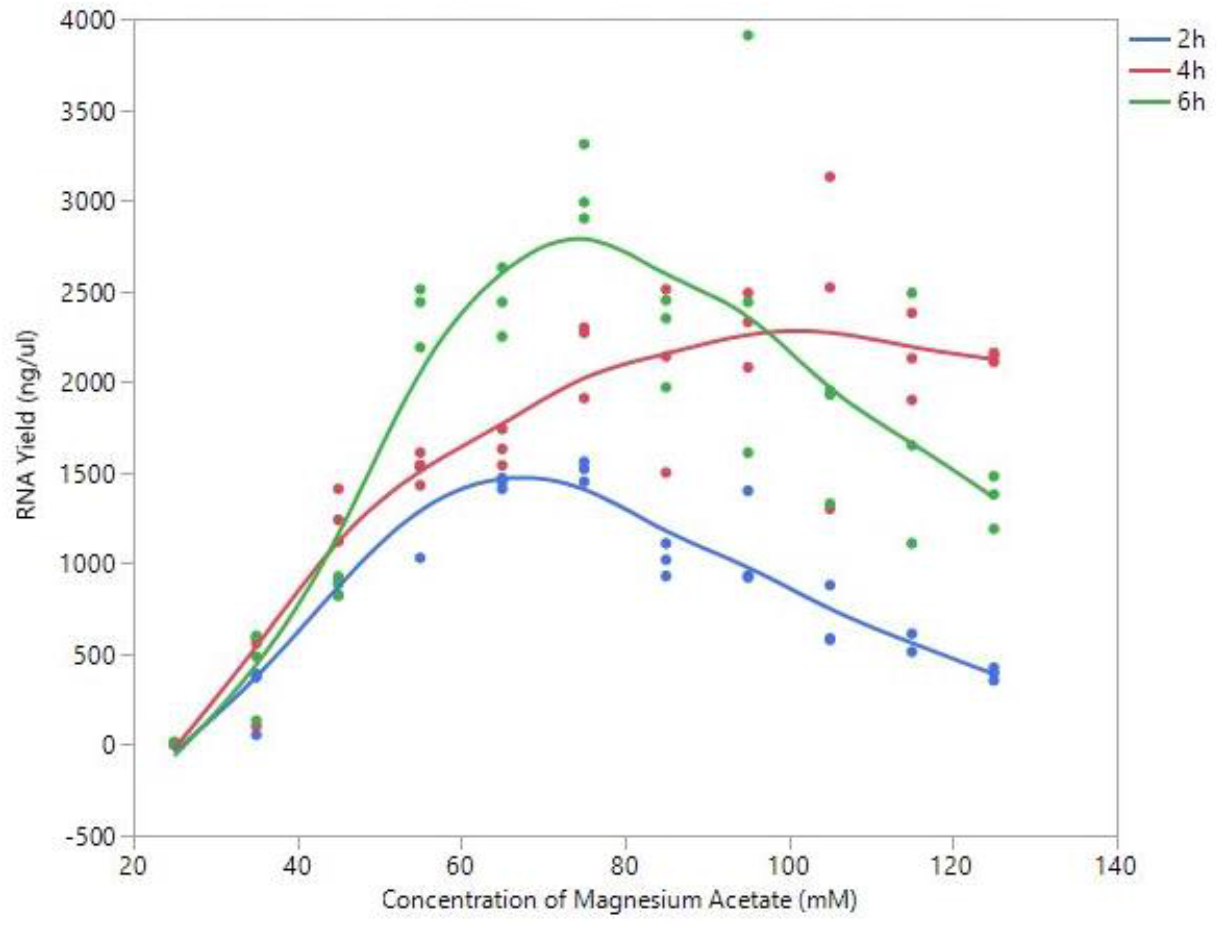
The impact of MgOAc_2_ on saRNA synthesis through IVT at different timepoints. Various IVT reactions were set up with different concentrations of MgOAc_2_ and the RNA yields were measured at 2, 4 and 6 h. Each MgOAc_2_ concentration varied was done in triplicates.

### RNA yield is proportional to T7 RNA polymerase concentration at all timepoints

Because many commercial kits recommend using 100U of T7 polymerase for IVT, we next wanted to explore whether lowering the T7 RNA polymerase concentration would yield the same RNA activity compared to 100 U of T7 RNA polymerase while increasing the incubation time. Hence, the optimal condition for IVT was used while varying only the concentration of T7 RNA polymerase, ranging from 12.5 to 100 U. Again, RNA yields were measured at 2, 4 and 6 h. We observed that lowering the T7 RNA polymerase concentration hinders the IVT process for all timepoints, leading to lower yields of RNA (Figure 7).

**Figure 7.**
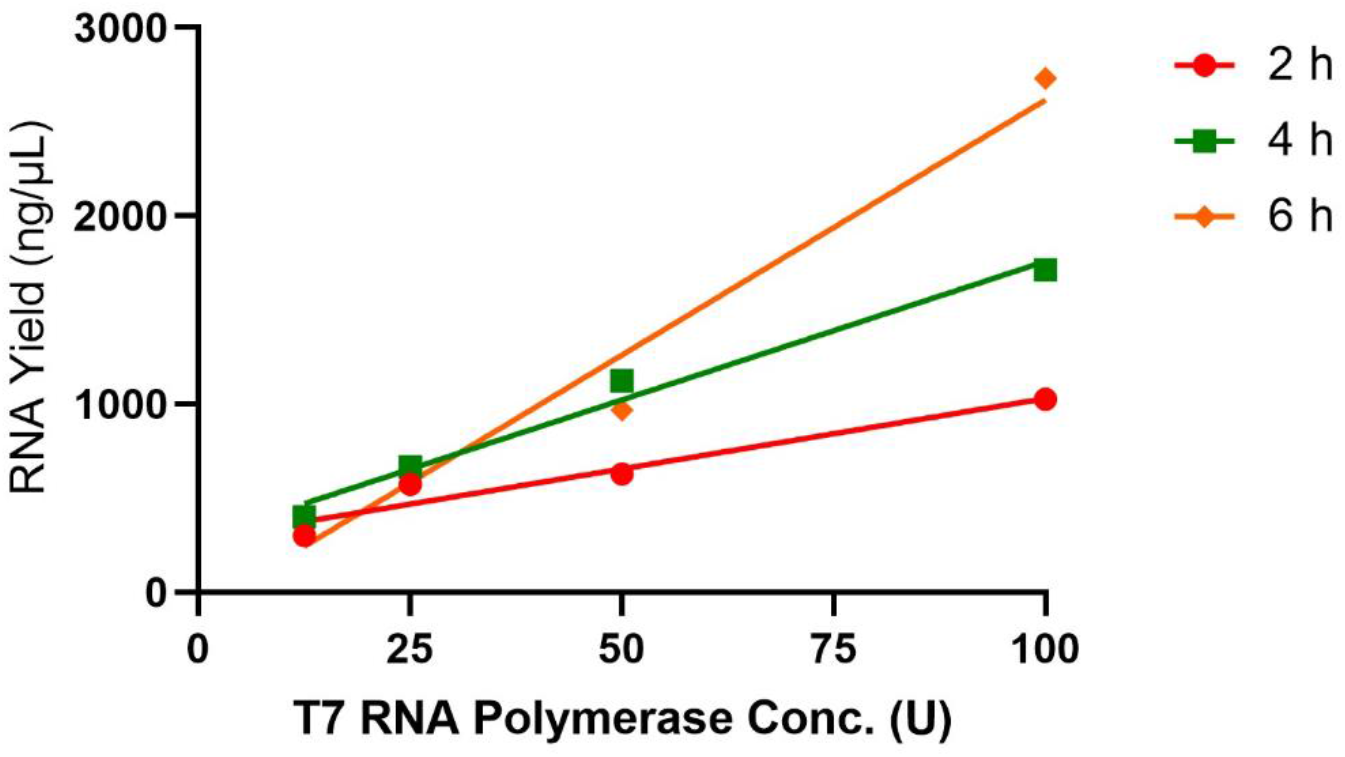
The effects of T7 RNA polymerase concentration on RNA yield at three different timepoints. Four different concentrations of T7 RNA polymerase were used for each IVT reaction and the RNA yields were measured at 2, 4 and 6 h. Each T7 RNA polymerase concentration varied was done in triplicates.

### Increasing the salt concentration during IVT process at different timepoints does not affect final RNA concentration

Because the binding efficiency of T7 polymerase is dependent on the ionic strength and the low-ionic-strength kinetics of RNA synthesis have been shown to plateau over time^22^, we then wanted to investigate whether increasing the salt concentration in the IVT reactions at different timepoints of the process might lead to higher RNA yields. Each reaction started off with 10 mM NaOAc and after incubating the reactions for 10, 30 and 60 min, various concentrations of NaOAc were added to the reaction to make a final concentration of 0.02, 0.1 and 0.5 M. Interestingly, the addition of different NaOAc concentrations yielded lower RNA concentrations than the reactions that had no increase in salt concentration (Figure 8), demonstrating that increasing the salt concentration during IVT process does not facilitate higher RNA yields.

**Figure 8.**
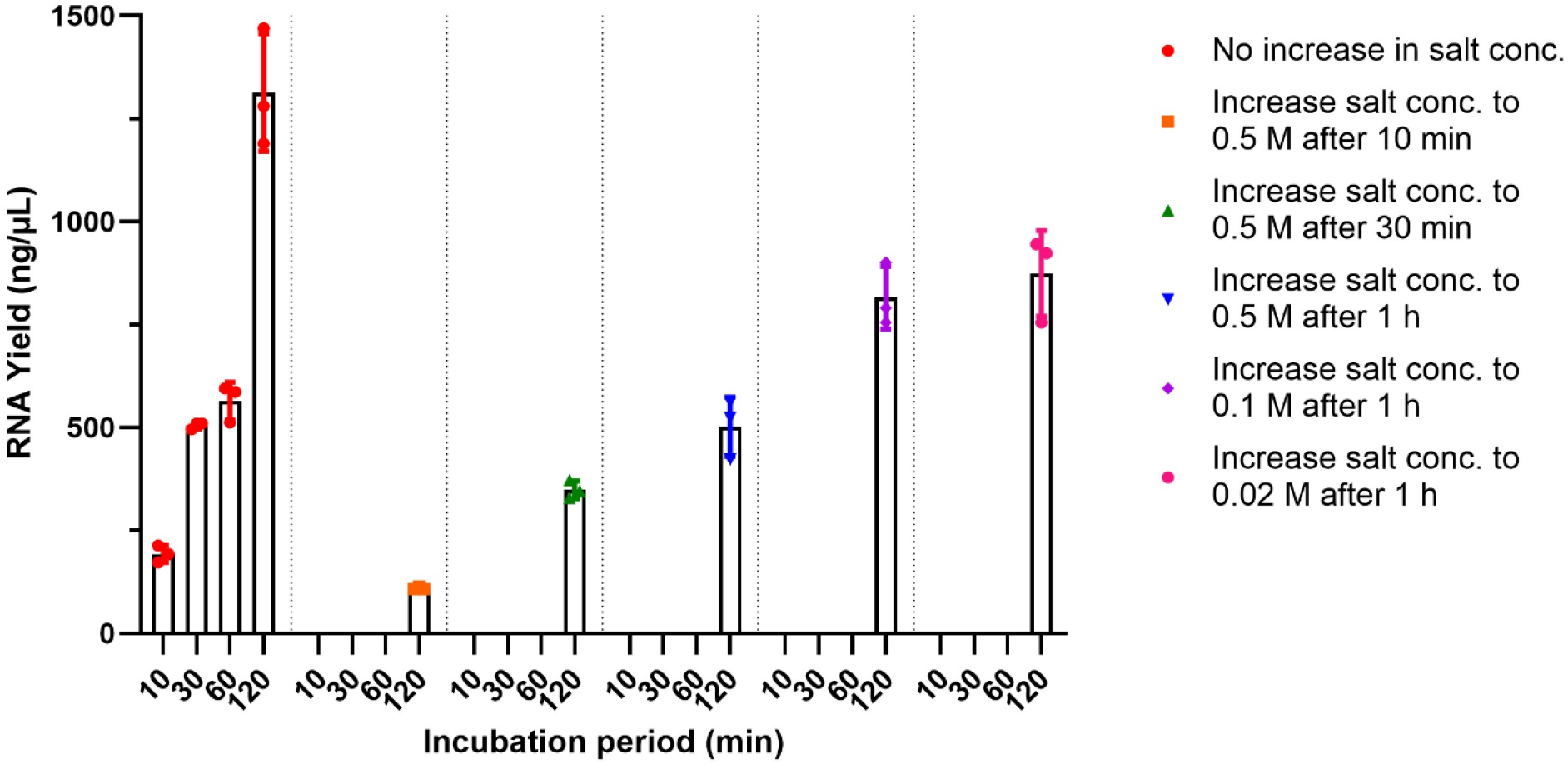
The effects of adding various NaOAc concentrations at different timepoints after the initiation of IVT on RNA yield. Different concentrations of NaOAc were added at either 10 min, 30 min or 1 h after IVT was initiated and the RNA yields of these reactions were measured at 2 h. The RNA yields of the reactions with no addition of sodium acetate were measured at 10 min, 30 min, 1 h and 2 h as a control. All conditions were done in triplicates.

### An optimal combination of pyrophosphatase, spermidine, DMSO, betaine and tween 20 is needed in order to produce high RNA yields

Next, we wanted to incorporate pyrophosphatase, spermidine, DMSO, betaine and a surfactant: either Triton X-100 or Tween 20; in order to see the interactions between each of these components in IVT. However, titrating these components individually would not allow us to determine these interactions. Therefore, we used a definitive screening design (DSD) of experiment where we took three different concentrations (a low, medium, and high point for each variable) and perform a minimum number of reactions to screen for effects and interactions, including quadratic effects. This allowed us to determine any curvature in the relationships between effects and estimate the local maximum. The concentrations used for each component are shown in Table 6. All RNA yields were measured at 4 and 6 h. Because the previous full three-level factorial DoE showed that 85 mM MgOAc_2_ worked best for a 4 h incubation period, we used 85 mM for this DoE. A Fit Definitive Screening platform was used to analyze the data in order to identify the main effects of IVT by using an Effective Model Selection for DSDs approach.

**Table 6.**
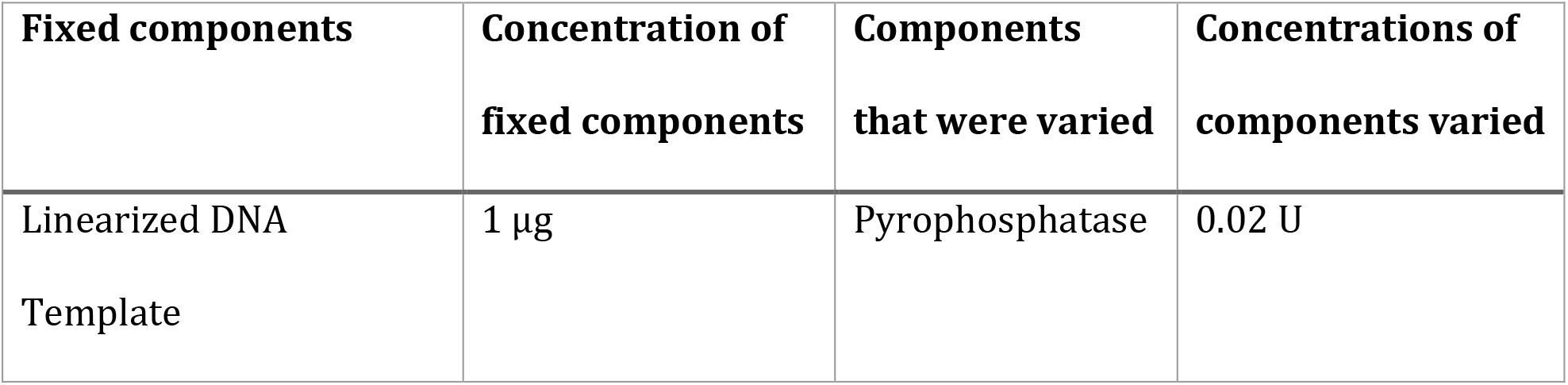
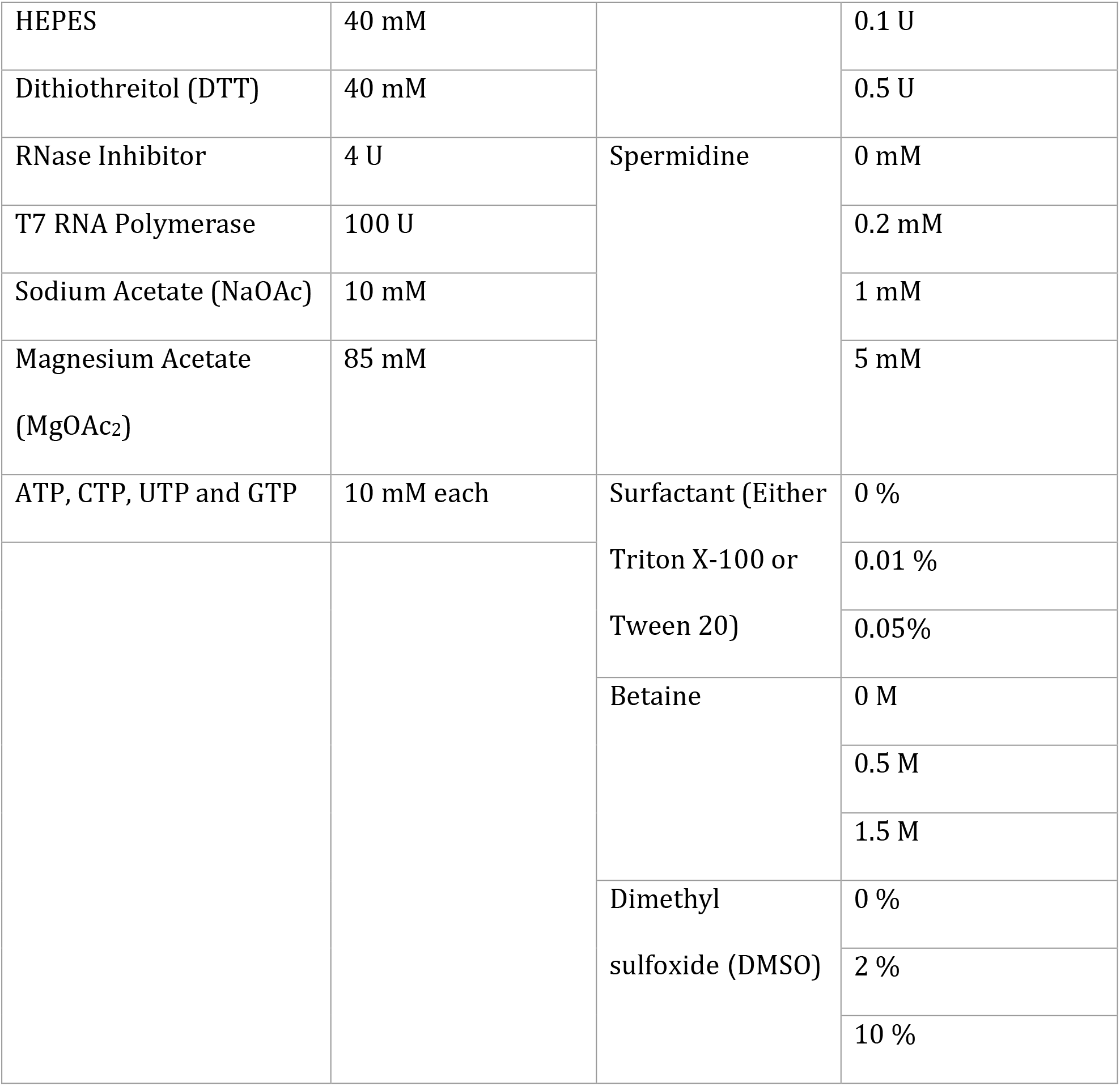
The components used for a definitive screening DoE.

We observed that high yields of RNA were caused by a variation of an optimal combination of each component. The major effect estimates showed that the spermidine, betaine and surfactant have the most impact on IVT with the surfactant being most significant (p= 0.0327) (Table 7). Interestingly, Tween 20 was shown to be more effective for increasing yield than Triton X-100 (Figure 9). However, pyrophosphatase and DMSO were shown to have a non-significant impact on IVT yield. In addition, for most of the components, high RNA yield is achieved only by using the medium concentration of each component rather than using the highest concentration, which suggests that too much of each component can result in a level that hinders saRNA production during IVT (Figure 9).

**Table 7.** The components that had major effects on IVT in a definitive screening DoE. Prob > |t| = p-value of the test. *A Prob > |t| value of <0.05 is considered significant.

| **Main Effect Estimates** |  |  |  |  |
| --- | --- | --- | --- | --- |
| **<u>Term</u>** | **<u>Estimate</u>** | **<u>Std Error</u>** | **<u>t Ratio</u>** | **<u>Prob > \|t\|</u>** |
| Spermidine | -491.3 | 211.58 | -2.322 | 0.0593 |
| Surfactant | 584.79 | 211.58 | 2.674 | 0.0327* |
| Betaine | 439.98 | 211.58 | 2.0795 | 0.0828 |

**Figure 9.**
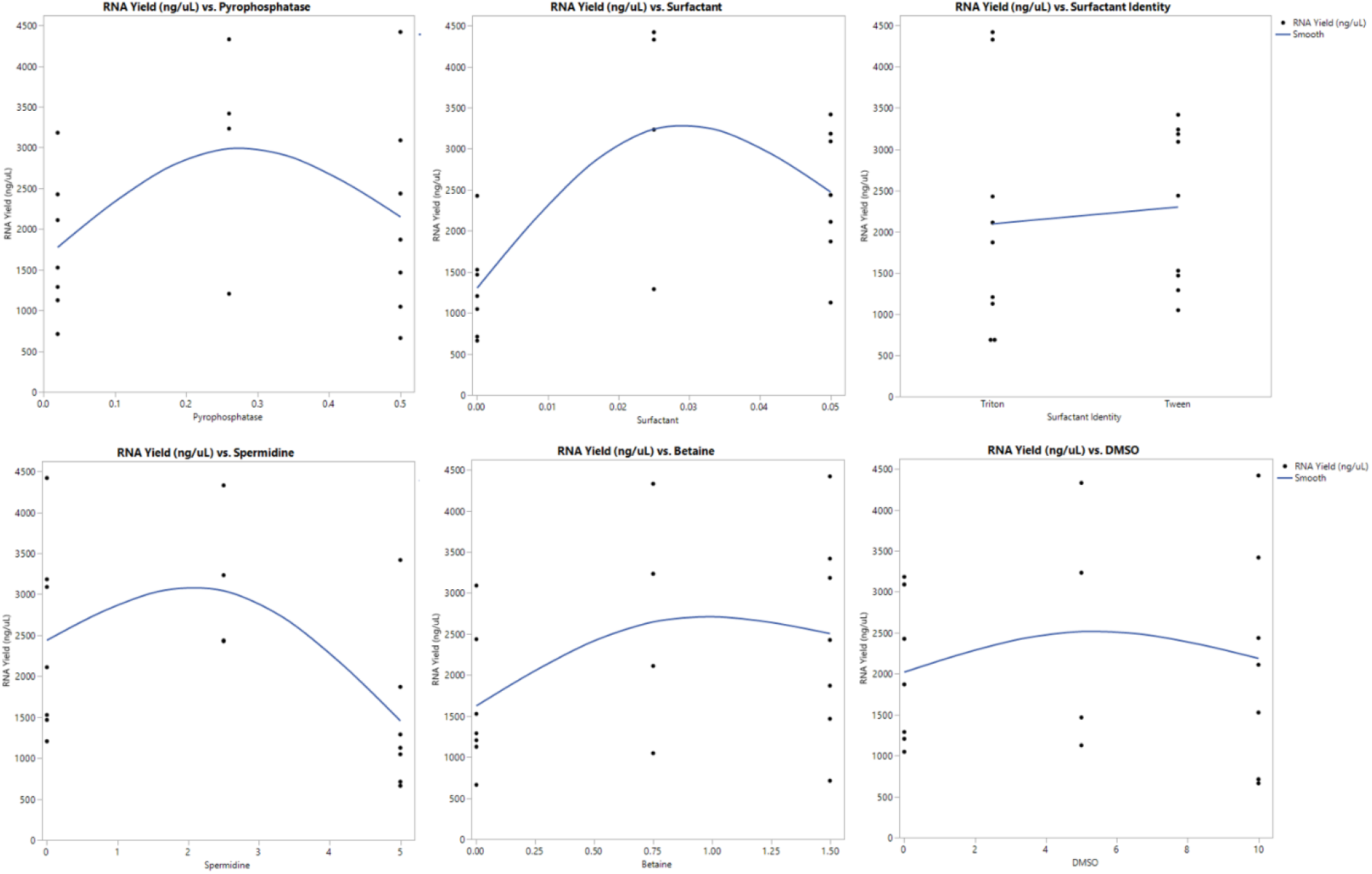
A definitive screening DoE analysis of saRNA production through IVT to observe the importance of pyrophosphatase, spermidine, surfactant (either Triton X-100 or Tween 20), betaine and DMSO on RNA yield. Each component varied was plotted against RNA yield to see their individual impact on RNA yield.

A color map of correlations also showed large absolute correlations between each effect, indicating that not only one effect is responsible for achieving high RNA yields but rather a combination of all components at their optimal concentration (Figure 10).

**Figure 10.**
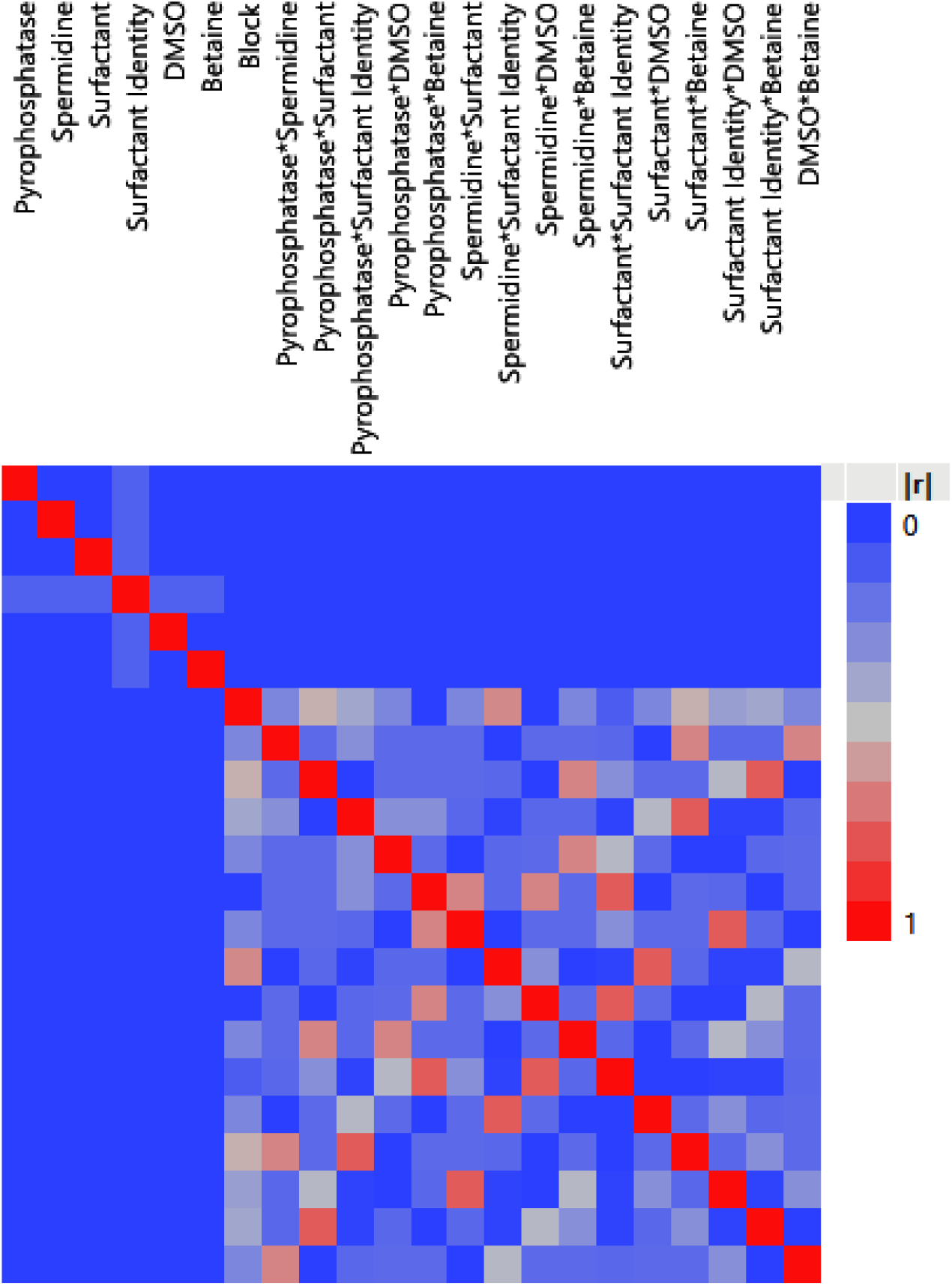
A color map on correlations between each component based on a definitive screening DoE analysis. The blue squares represent small correlations between each component. The gray squares represent correlations between components with quadratic effects. The red squares represent high correlations. |r| represents the absolute value of the correlation between components. Components with large absolute correlations indicate an increase in the standard errors of estimates.

### Inclusion of pyrophosphatase does not enhance the quantity of RNA yield

Because many IVT protocols include pyrophosphatase for catalyzing the hydrolysis of any pyrophosphate byproduct which can limit polymerization rate and synthesize shorter RNA molecules^23^, we wanted to explore whether pyrophosphatase was necessary in generating high RNA concentrations. We compared our two current best conditions for IVT with and without pyrophosphatase and measured RNA concentrations after 2 and 4 h incubation. Both conditions yielded similar concentrations at each timepoint (Figure 11A). However, analysis of RNA quality by gel electrophoresis indicated that the condition without pyrophosphatase showed a slightly better quality of RNA for both the 2 and 4 h timepoints compared to the condition with pyrophosphatase (Figure 11B). This suggests that pyrophosphatase provided no benefit in generating high RNA yield under these optimized conditions.

**Figure 11.**
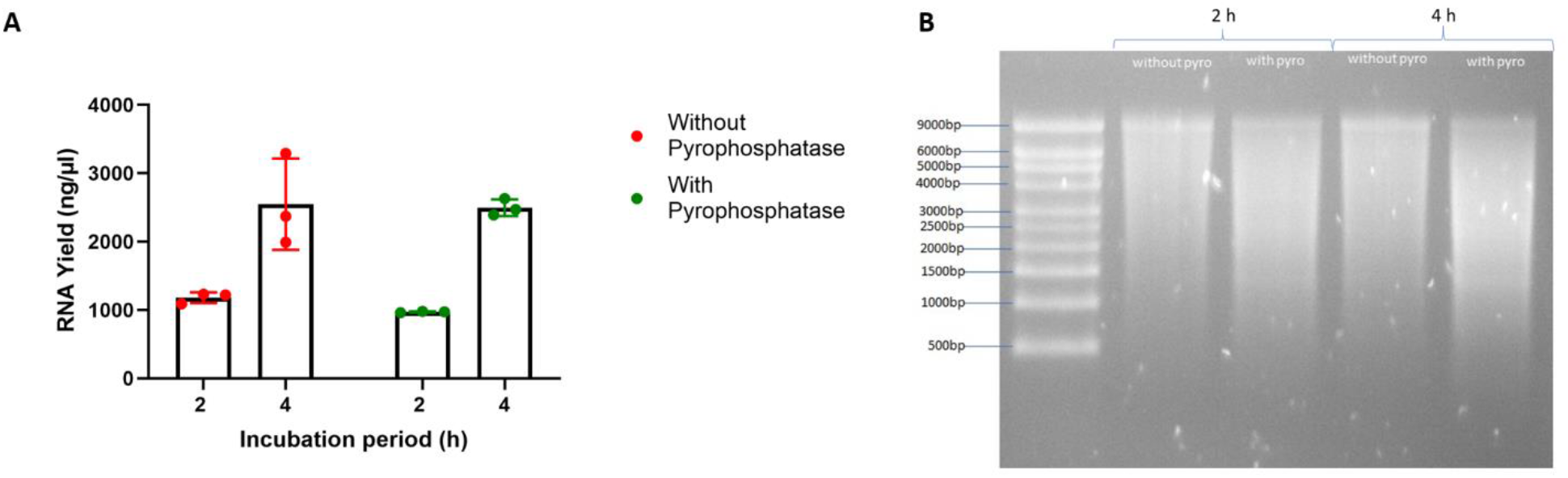
Comparison of the current best conditions with and without pyrophosphatase. (A) After conducting all DoEs, the optimal conditions with and without pyrophosphatase were set up for IVT to compare the RNA yield at 2 and 4 h timepoint. (B) The RNA replicons synthesized at 2 and 4 h were then purified and analzyed on a denaturing gel for quality control.

### The optimal conditions of IVT can be used to synthesize saRNAs encoding for different gene of interests with good protein expression *in vitro*

We then wanted to confirm that our optimal IVT condition could be used to synthesize different sizes of RNA. Thus, we prepared four other pDNA templates for making saRNA encoding different GOIs that included enhanced green fluorescent protein (eGFP) (8.5 kb), multi-drug resistance-1 (MDR1) (11.5 kb) and multi-drug resistance-1 with parainfluenza virus 5 (MDR1-PIV5) (12.4 kb). In addition, we used a pDNA template encoding for fLuc to synthesize mRNA fLuc (1.8 kb). The sizes of each RNA species ranged from 1.8 to 12.4 kb. Our optimized IVT condition (Table 8) generated high yields of good quantity and quality RNA irrespective of size (Figure 12A and B). The total mass yielded for VEEV-fLuc, VEEV-eGFP, VEEV-MDR1, VEEV-MDR1-PIV5, and mRNA fLuc were 118, 150, 142, 169 and 41 μg, respectively; while the average moles of VEEV-fLuc, VEEV-eGFP, VEEV-MDR1, VEEV-MDR1-PIV5, and mRNA fLuc were 38.94, 54.77, 38.32, 42.48 and 70.28 pmol, respectively. Lastly, we transfected these RNAs into HEK293T.17 cells and harvested the cells 48 h post transfection in order to confirm the protein expression from each GOI for saRNAs and for the fLuc mRNA. We observed appropriate protein expression from each of the saRNA and the fLuc mRNA (Figure 12C).

**Table 8.** The final optmized components for IVT.

| **Components** | **Concentration of each component** |
| --- | --- |
| Linearized DNA Template | 1 µg |
| HEPES | 40 mM |
| Dithiothreitol (DTT) | 40 mM |
| RNase Inhibitor | 4 U |
| T7 RNA Polymerase | 100 U |
| Sodium Acetate (NaOAc) | 10 mM |
| Magnesium Acetate (MgOAc<sub>2</sub>) | 75 mM |
| ATP, CTP, UTP and GTP | 10 mM each |
| Spermidine | 0.2 mM |

**Figure 12.**
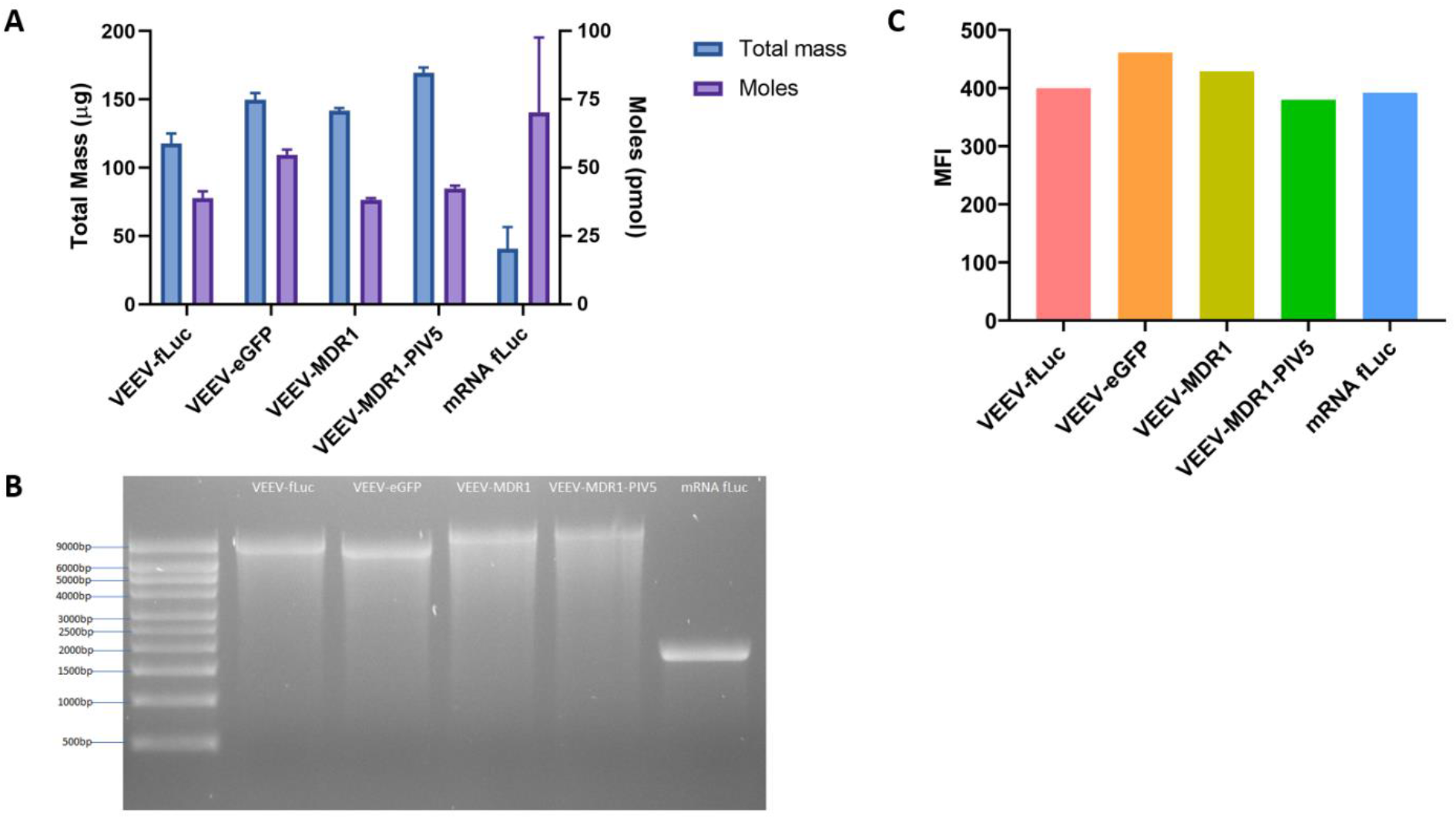
Comparison of the current best condition without pyrophosphatase with different genes of interest. (A) The current best condition without pyrophosphatase was used to synthesize RNA replicons containing different GOIs as well as mRNA fLuc through IVT for 2 h and the RNA yield was measured using the Qubit. Total mass and moles of each RNA were quantified. (B) RNAs were then purified and ran on a denaturing gel for quality control. (C) Each RNA was transfected into a 6-well plate of HEK293T.17 cells. At 48 h post transfection, cells were harvested and stained with appropriate antibodies to observe protein expression of each GOI by flow cytometry, in which the expression level is represented as the median fluorescence intensity (MFI).

## Discussion

The aim of this study was to utilize a DoE approach to optimize the IVT process for long saRNAs so that maximal RNA yield can be achieved. Here, we found that magnesium plays the most important role in IVT, especially its interaction with NTPs where the right balance of these two components is needed to yield high RNA concentrations. In addition, acetate ions were more effective for IVT compared to chloride ions with IVT being most effective at 75mM MgOAc_2_ and 10 mM of each NTP. Maintenance of ionic strength through the addition of NaOAc at different timepoints of the IVT process did not enable the production of high RNA yields. We also found that lowering T7 polymerase hinders IVT process and that pyrophosphatase is not significantly required for IVT in order to yield maximal RNA concentrations. Finally, we observed that the optimized condition for IVT could be used to produce a range of RNA, from 1.8 to 12.4 kb and that each type of RNA was functional and thus able to express proteins once transfected into cells.

Despite the T7 polymerase being the most essential component in order to initiate IVT^24^, other factors influence optimal IVT productivity. Magnesium ions are known to influence the catalytic activity of T7 polymerase^25^. However, the ratio of magnesium to NTPs is the critical parameter influencing efficient IVT. We observed that a combination of 10 mM of each NTP with 75 mM of magnesium anion produced optimal IVT RNA yield, where 5 mM of each NTP is half as good and 20 mM of each NTP being too much. This means the optimal molecular ratio between total NTPs and magnesium concentration is 1: 1.1875 (M/M). The relationship between NTPs and magnesium found in this study corresponds to previous research whereby if a low concentration of NTPs was used, it limits RNA production but high concentrations of NTPs become inhibitory^26^. One explanation for this phenomenon is that when hydrogen ions are released from the NTP during the formation of the magnesium-NTP complex, the pH of the reaction is reduced which blocks the binding between the T7 RNA polymerase and the DNA^27^. This indicates that the higher the NTP and magnesium concentration, the more hydrogen ions are released and the fall in pH is greater, subsequently inhibiting enzyme activity. Therefore, the optimal amount of NTPs must be used in relation to the amount of magnesium added initially in order to synthesize high yields of RNA.

Previous research has shown that acetate anions are more effective during IVT compared to chloride anions. Activity of the T7 RNA polymerase is more sensitive to the chloride anions when the polymerase binds to the T7 promoter, in which the chloride anion competes with the DNA to bind to specific protein sites^28^. This causes a decrease in transcription efficiency^29^. Hence, when we performed a DoE to explore the interactions between sodium, magnesium and NTPs, we also compared the differences between using chloride and acetate anions. Similar to previous studies of small RNA IVT, our data using long RNA also indicate that the ion type is a significant factor in IVT with acetate anions being more effective than chloride anions.

Interestingly, while many protocols use pyrophosphatase for IVT^23, 30^, we found that pyrophosphatase was not necessarily required to yield high RNA concentrations. When the T7 polymerase binds to the DNA template, this complex binds to the magnesium-NTPs complex and pyrophosphate is released, in which this pyrophosphate byproduct has been shown to slow down IVT^27^. Hence pyrophosphatase is incorporated to eliminate any pyrophosphate byproducts. However, because the free magnesium ions precipitate pyrophosphate^27^, we hypothesize that pyrophosphatase had little impact in our DoEs because the magnesium levels were high enough to cope with the generated pyrophosphate levels.

Furthermore, previous research has found that RNA synthesis by DNA-dependent RNA polymerases like the T7 RNA polymerase is highly influenced by the ionic strength^22, 31^. Fuchs et al. found that increasing the ionic strength 10 min after the initiation of IVT to a high ionic strength allowed the IVT reaction to run maximally for 6 h^31^. An explanation for this was that under low ionic strength, the T7 polymerase is only able to make a single copy of the transcript and only has the ability to re-initiate the reaction to make additional copies when the ionic strength is high. However, too high of an ionic strength might interfere with the binding activity of the T7 polymerase to the DNA template^32^. We observed no benefit from the addition of sodium acetate during our IVT reactions. This likely reflects that the ionic strength was already optimal in our IVT setup.

Since many IVT protocols have been optimized for shorter mRNAs, the application of the DoE approach proved instructive to understanding the interactions between each reaction component to produce maximal yields of long RNA. Further benefits could be gained by determining the optimal conditions for co-capping of saRNA during IVT. However, our optimized IVT reaction conditions that generate high yields of saRNA can be used to produce large quantities of saRNA for research and development purposes and have the potential to be scaled up for GMP production of saRNA vaccines and biotherapeutics.

## Abbreviations used

saRNA: self-amplifying RNA;
IVT: in vitro transcription;
DoE: design of experiment;
pDNA: plasmid DNA;
GOI: gene of interest;
NTP: nucleoside triphosphate;
VEEV: Venezuelan Equine Encephalitis Virus;
firefly luciferase: fLuc

## Data Availability

Raw data is available upon request through Imperial College London.

## Author Contributions

KS, AKB, PFM and RJS conceptualized and designed the studies. KS performed the *in vitro* experiments. KS and AKB analyzed the data. KS wrote the manuscript with constructive feedback and editing from AKB, PFM and RJS.

## Acknowledgements

We acknowledge the Dormeur Investment Service Ltd. For providing funds to purchase equipment used in these studies.

